# *MECP2* Mutations Rewire Human ESC Fate and Bias Cortical Lineage Commitment

**DOI:** 10.1101/2025.09.30.679576

**Authors:** Marion Guillon, Margaux Brin, Elodie Gabet, Justine Gromaire, Mathéa Bernard, Laetitia Laurent, Théo Rabin, Laila Asali, Yi Liu, Anthony Flamier

## Abstract

Rett syndrome arises from loss-of-function mutations in the X-linked chromatin regulator *MECP2*, yet the earliest molecular derailments in human development remain poorly defined. Using isogenic hESC models carrying three patient-derived *MECP2* mutations, we followed the transcriptome from pluripotency through neuro-ectoderm, neural stem, and neural progenitor stages and into three-month cerebral organoids. Stage dominated transcriptional variance, but mutants shared a secondary program enriched for synaptic-membrane and extracellular-matrix genes. Single-cell profiling revealed a naïve-like, hyper-proliferative state marked by up-regulation of *ZFP42* at ESC stage. Strikingly, *EMX1*, a cortical radial-glia determinant, was consistently suppressed from the earliest stage onward, and cerebral organoids subsequently generated fewer excitatory neurons in favour of inhibitory and glial lineages. These data chart a continuous developmental trajectory for *MECP2*-mutant human cells and nominate ZFP42 and EMX1 dysregulation as tractable entry points for dissecting Rett pathogenesis.

## Introduction

Rett syndrome (RTT) is a severe X-linked neurodevelopmental disorder that affects ≈1 in 10,000 girls worldwide and, in rare surviving males, presents with an even more severe encephalopathy.^1,2^ After a seemingly normal first 6–18 months, infants enter a protracted period of neuro-regression marked by loss of purposeful hand use and speech, seizures, gait ataxia, autistic features, and severe intellectual disability.^1–8^ Structural MRI and post-mortem studies consistently show reduced total brain volume, cortical thinning, simplified dendritic arbors, and a shift in excitatory– inhibitory balance, suggesting that pathogenic processes likely begin early corticogenesis and extend into post-natal synaptic maturation.^9–13^

More than 95 % of classic RTT cases arise from de-novo loss-of-function mutations in *MECP2*, which encodes methyl-CpG-binding protein 2 (MECP2).^14^ Historically, MECP2 was viewed as a canonical methyl-DNA reader that blankets the neuronal genome and recruits co-repressor complexes to silence transcription.^15^ However, recent molecular studies on adult mouse cortex and human embryonic stem cells (ESCs) derived neurons refine this view: MECP2 binds preferentially to discrete enhancer-like elements, termed MECP2-binding hotspots (MBHs), and does so largely independently of local CpG methylation.^2,16–19^ Clusters of MBHs cooperate to dampen transcription of genes enriched for neuronal functions, revealing an intragenic, methylation-independent mode of repression that likely complements the classical methylation-dependent mechanism.^16,17^ We and others also found that MECP2 binds both unmethylated and methylated cytosines to act both as transcriptional repressor (through its NcoR domain) and activator (through RNA polymerase II recruitment) in neurons.^17,18,20–22^ However, the mechanistic context of these findings remains unclear in early development. Mammalian ESCs and blastocyst-stage embryos are globally hypomethylated, a state long thought to preclude meaningful MECP2 engagement and thus to spare pluripotent cells from RTT-associated lesions.^23^ The discovery that MECP2 can bind hypomethylated regions to regulate transcription now raises the possibility that *MECP2* deficiency could perturb transcriptional programs as early as the hypomethylated ESC stage, well before neurons are specified.^2,17^ Determining whether such early dysregulation contributes to later cortical deficits is therefore a central unanswered question that we address in this study.

To interrogate this possibility in a continuous human model, we used isogenic human male hESC lines harboring three recurrent RTT mutations (R133C, R168X, R270X)^24^ and performed longitudinal profiling across four matched developmental milestones — pluripotent ESC, neuro-ectoderm (NE), neural stem cell (NSC) and neural progenitor cell (NPC) — and in long-term cerebral organoids. We further used RTT patient-derived induced pluripotent stem cell (iPSC) lines to validate our observations. By integrating bulk and single-cell RNA sequencing with gene-set enrichment and machine-learning classifiers, we uncovered a stage-persistent, MECP2-dependent transcriptional program that elevates naïve stem cell markers, accelerates ESC proliferation, increases synaptic-membrane pathways during neural induction, and suppresses the cortical radial-glia determinant EMX1, culminating in a quantitative deficit of excitatory neurons. These data reveal discrete molecular lesions that precede overt neuronal dysfunction and provide new developmental entry points for mechanistic dissection in RTT.

## Results

### Stage-resolved transcriptomics reveals early, genotype-specific disruptions in *MECP2*-mutant differentiation

To test whether MECP2 loss perturbs human neurodevelopment from its earliest hypomethylated state, we profiled three previously described CRISPR-edited, isogenic hESC lines carrying common RTT mutations (R133C, R168X, R270X) alongside WT controls as they were sequentially coaxed through ESC (Day 0), NE (Day 3), NSC (Day 7) and NPC (Day 21) stages (Figure 1A).^17^ As expected, the pluripotency factors plunged after the ESC stage, whereas neuronal markers became detectable at NSC and rose thereafter, confirming efficient lineage induction (Figure 1A and S1A). *MECP2* itself remained low but readily measurable at ESCs (5–10 TPM), corroborating published observations (Figure 1A).^18^ *MECP2* increased modestly at NE/NSC and surged at NPC (Figure 1A). This suggests that residual embryonic expression could already influence transcriptional programs. Principal-component analysis showed that PC1 (59 % variance) stratified samples by developmental stage, validating the differentiation axis, whereas PC2 (13 %) captured mutation-specific variance that emerged progressively: R270X diverged from WT at the ESC stage, R168X detached by NE, and all mutants formed distinct genotype clusters by NSC/NPC (Figure 1B). GO enrichment revealed significant dysregulation beginning at ESCs, most strikingly in R270X where synaptic-membrane genes were already mis-expressed (Figure 1C). NSC samples displayed enrichment for “regulation of nervous-system development,” and NPCs showed broad perturbation of synaptic and ion-channel terms (Figure 1C). We observed common gene dysregulations across genotypes and stages (Figure 1C and S1B). Two transcripts were consistently altered in every genotype at every stage: *S100A6*, a calcium-binding protein implicated in cytoskeletal dynamics and stem-cell proliferation, and *SLC17A7*/*VGLUT1*, the principal vesicular glutamate transporter essential for excitatory neurotransmission (Figure 1C).^25–27^ Early dysregulation of *S100A6* suggests cytoskeletal or proliferative defects could precede neurogenesis, whereas persistent mis-expression of *SLC17A7* foreshadows later synaptic dysfunction.^27–29^ Collectively, these findings suggest that *MECP2* mutations impose transcriptomic disturbances as early as the blastocyst-like ESC stage, raising the possibility that altered lineage trajectories, not only neuronal maturation defects, contribute to RTT pathogenesis.

**Figure 1.**
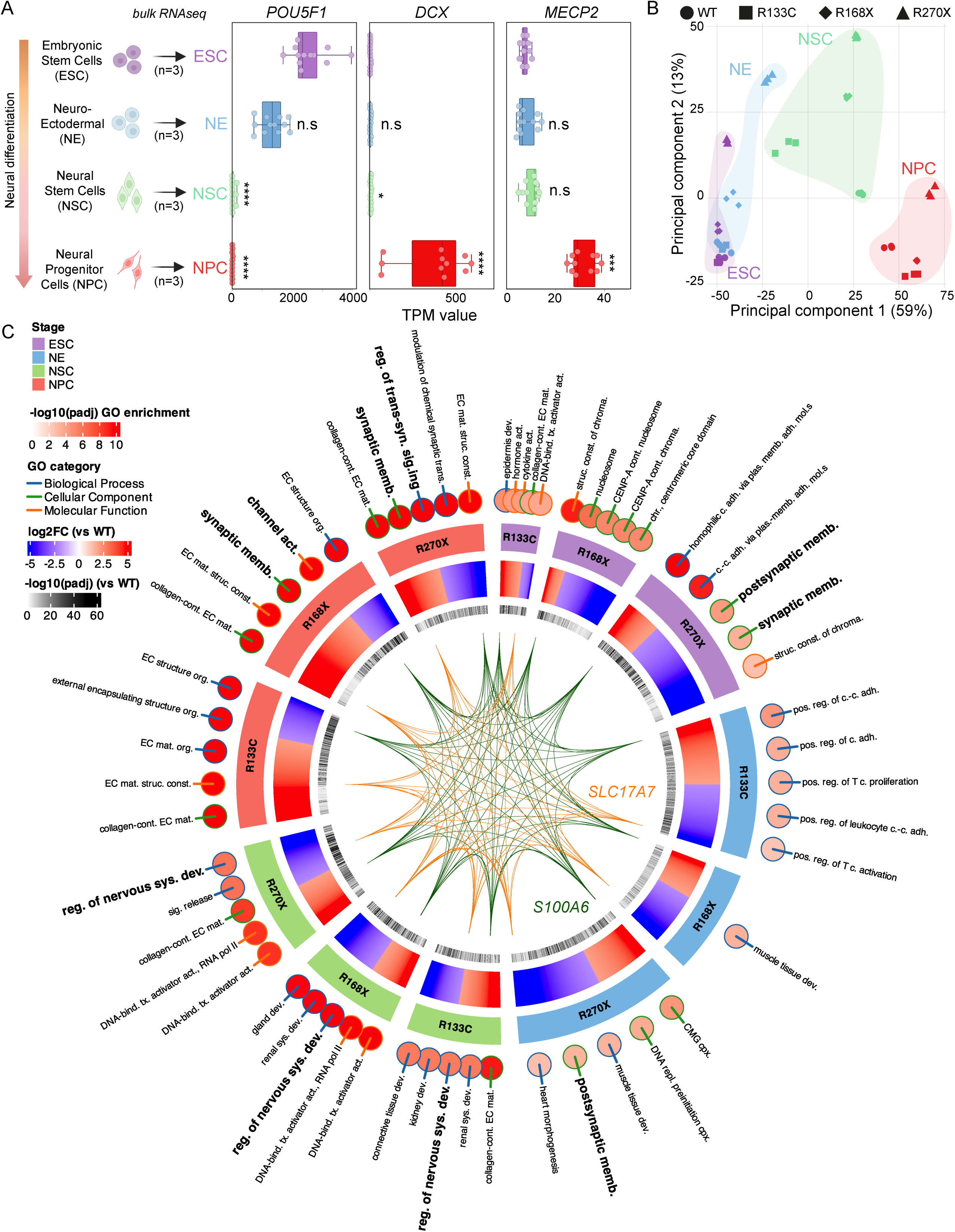
Stage-resolved transcriptomics reveal early gene-network disruption in *MECP2*-mutant hESC differentiation. **A.** Expression dynamics of key developmental markers across the four matched stages analyzed on 4 RTT hES cell lines (WT, R133C, R168X et R270X). Box-and-scatter plots show transcripts per million (TPM; n = 3 biological replicates per stage and genotype) for the pluripotency factor *POU5F1*/OCT4, the early neuronal marker *DCX*, and *MECP2* itself. N=3; Tukey-adjusted post-hoc significance relative to the ESC stage (\**P* < 0.05, *** *P* < 0.001, **** *P* < 0.0001). **B.** Principal-component analysis of the complete bulk RNA-seq dataset. **C.** Circular multi-layer GO enrichment map integrating all three mutants. Outer colored dots show top 5 GO terms significantly enriched at each stage (ESC, NE, NSC, NPC) and for each genotype versus WT (neural GO terms in bold); color scale encodes log2 fold-change versus WT (red, up-regulated; blue, down-regulated). Inner ring indicates –log10 padj of differential gene expression. The centre chord diagram links the two transcripts (*S100A6*, green chords; *SLC17A7*, orange chords) that are consistently dysregulated in every genotype at every stage.

### Machine-learning models accurately decode developmental stage and highlight early neuronal mis-timing in *MECP2* mutants

We asked whether *MECP2* mutations create transcriptomic shifts large enough for detection by machine-learning and, if so, which genes drive those shifts. To this end we analysed our bulk RNA sequencing matrix with three complementary algorithms: unsupervised k-means clustering served for pattern discovery; a supervised random-forest ensemble offered interpretable prediction; a feed-forward neural network provided high-capacity validation. K-means partitioned genes into six clusters (Figure 2A and S2A-B). Cluster 6 showed a steady increase in expression from ESC to NE, NSC and NPC (Figure 2B and S2B). Notably, this trajectory was the most significantly altered in the three mutant lines (Figure 2A-B). Gene-Ontology analysis linked this cluster to axonogenesis and regulation of nervous-system development, implying that neuronal programs are engaged unusually early in *MECP2*-deficient pluripotent cells (Figure 2C). The random forest, trained on labelled samples, achieved more than 95 % cross-validated accuracy (Figure 2D). Feature-importance scores highlighted ten transcripts that dominate stage discrimination (Figure 2E). For example, *GABRE,* encoding the ε-subunit of the GABA-A receptor shaping inhibitory tone was elevated in mutants at ESC and NE but normalised or fell below wild-type at NSC and NPC (Figure 2F).^30,31^ *PDCD2L*, with a similar profile, plays a role in neurodevelopment, particularly in programmed cell death (PCD) and ribosomal biogenesis (Figure 2F).^32^ PCD is essential for stem cell maintenance and neurodevelopment, as it involves the elimination of a subset of neurons to refine neural circuits.^33^ Finally, *NSMCE4A*, the most important gene of this classifier, is part of the SMC5/6 complex which is key for chromosome architecture and genomic stability during neurogenesis.^34,35^ The gene expression profile mirrors Cluster 6 and suggests premature activation followed by later exhaustion of inhibitory-synapse genes (Figure 2B). A feed-forward neural network trained on the 2,000 most variable genes reproduced stage calls with all WT samples on the diagonal of true versus predicted stage (Figure 2G). This model suggests that the R270X line exhibited the most divergent alignment from the WT trajectory, particularly in the NE stage. Convergence across k-means, random forest and neural network indicates that developmental stage still dominates variance, yet *MECP2* mutations superimpose an early surge and later collapse of neuronal and synaptic transcripts. This temporal mis-alignment helps automated models flag mutant ESCs and underscores the need to dissect transcriptional disruption at the blastocyst-equivalent stage.

**Figure 2.**
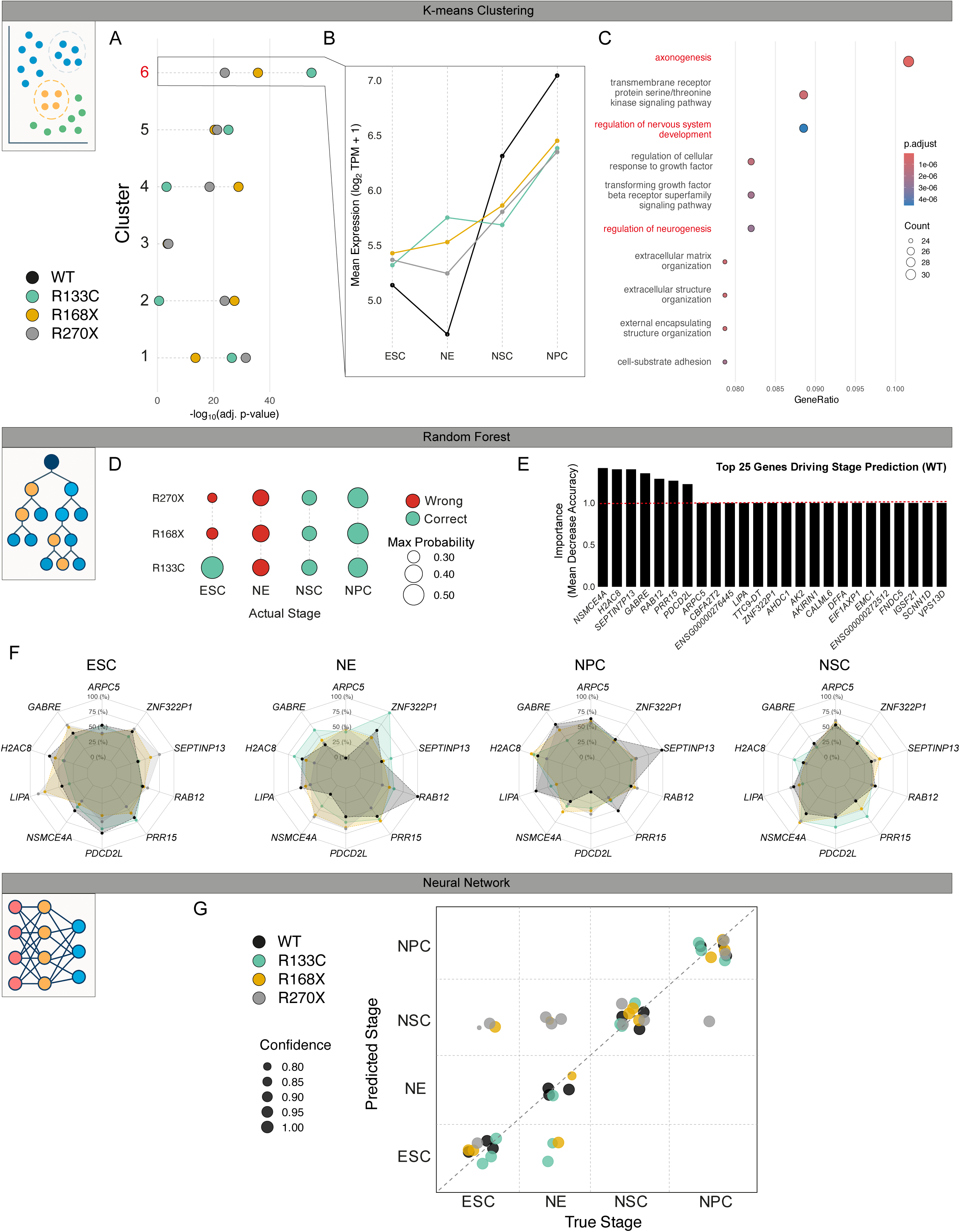
Machine-learning classifiers expose premature neuronal program activation in *MECP2* mutants. **A.** Adjusted p-values (–log₁₀ scale) for the six gene clusters identified by K-means (left margin indicates cluster number). **B.** Mean expression profile (log₂ TPM + 1) of the genes represented in cluster 6 across the four sampled stages and the four genotypes. Genotypes are color-coded as in the inset legend. **C.** Dot plot of GO terms enriched in Cluster 6. Dot diameter corresponds to the number of genes annotated to the term; color encodes Benjamini–Hochberg adjusted p-value. **D.** Confusion matrix displaying random-forest stage predictions for each genotype. Circle size denotes the classifier’s maximum posterior probability; color indicates correct or incorrect assignment. **E.** Bar chart of the top 25 genes ranked by mean decrease in accuracy when permuted, obtained from the model trained on WT samples. **F.** Radar plots of the ten most informative transcripts (panel E) showing their median expression at each stage for WT and the three *MECP2*-mutant lines. **G.** Scatter plot comparing predicted versus true developmental stages for all samples. Point color denotes genotype; point size reflects the model’s confidence (posterior probability). The dashed diagonal marks perfect agreement.

### *MECP2* loss promotes a naïve-like transcriptional program and hyperproliferation in human ESCs

To determine whether the early transcriptional disruption at the blastocyst-equivalent stage arises from heterogeneous differentiation states, we performed single-cell RNA sequencing on undifferentiated hESCs carrying WT, R133C, R168X or R270X *MECP2* alleles. UMAP embedding revealed four genotype-specific clusters: WT and R133C cells co-localized, whereas R168X and R270X formed distinct groupings that recapitulate our bulk RNA sequencing observations (Figure 1B and 3A); the absence of subclusters within each genotype confirms homogeneity (Figure 3A). Expression of the core pluripotency factor *POU5F1* was uniform, yet truncating mutants (R168X and R270X) exhibited striking upregulation of the gene *ZFP42* (Figure 3B). *ZFP42*, also known as *REX1*, is a gene that encodes a zinc finger protein primarily expressed in undifferentiated stem cells.^36,37^ *ZFP42* is a marker of naive pluripotency in ESCs, an early, less differentiated state of stem cells, while primed pluripotency is a more developmentally advanced state.^38–41^ The pronounced upregulation of *ZFP42* in truncating mutants raises the possibility that these cell lines have acquired aberrant pluripotency states or compromised genomic stability.

**Figure 3.**
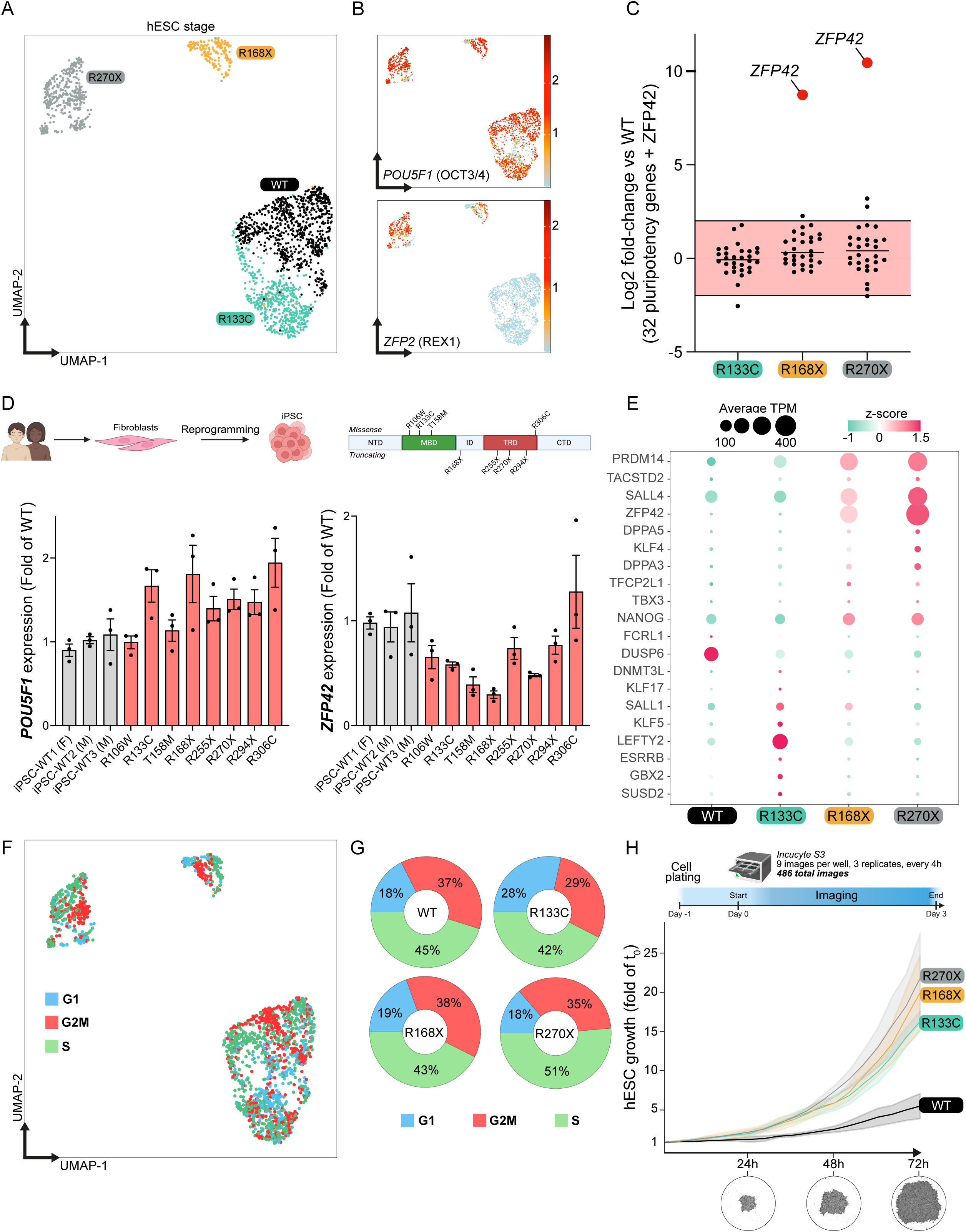
MECP2 loss drives a naïve-like transcriptional shift and hyperproliferation in human ESCs. **A.** UMAP projection of 2,925 single hESC transcriptomes colored by genotype (WT, R133C, R168X, R270X) at the pluripotent stage. **B.** Feature plots showing the per-cell log-normalized expression of the core pluripotency factor *POU5F1*/OCT4 (top) and the naïve-associated marker *ZFP42*/*REX1* (bottom) on the same UMAP embedding. **C.** Dot plot summarizing log₂ fold-change (mutant vs WT) for a panel of 32 canonical pluripotency genes (black dots); red dots highlight *ZFP42* values. Computed from bulk RNA sequencing matrix. **D.** Top, schematic of fibroblast reprogramming to iPSCs; bottom, bar graphs (mean ± s.e.m., n = 3) of *POU5F1* and *ZFP42* transcript levels in three control iPSC lines and eight Rett iPSC lines harboring the indicated *MECP2* alleles. (F: Female; M: Male). **E.** Dot plot displaying average expression (dot size) and z-score (color scale) of the 20 naïve-enriched stem-cell markers across WT and mutant hESCs. Computed from bulk RNA sequencing matrix. **F.** UMAP embedding from panel A with cells colored by inferred cell-cycle phase (G₁, S, G₂/M) using gene-signature scoring. **G.** Donut plots quantifying the proportion of cells in each cell cycle phase for WT and the three mutants. **H.** Live-cell imaging proliferation assay, growth curves of hESC colonies over 72 h recorded on an Incucyte S3 (9 images/well, N=3); inset photographs illustrate colony morphology at 24, 48 and 72 h.

Survey of 32 canonical pluripotency genes revealed no significant dysregulation, excluding a broad shift in pluripotency (Figure 3C). InferCNV analysis of our single-cell transcriptomes detected no copy number alterations, and whole-genome long read sequencing confirmed that all lines retained the WT genotype (Figures S3 and S4). Accordingly, the increased *ZFP42* expression in R168X and R270X hESCs may be indicative of a more naïve-like transcriptional state, rather than evidence of underlying pluripotency defects or genomic instability. We next analyzed three control iPSC lines alongside eight patient-derived Rett iPSC lines (Figure 3D). In our male model there is only one X chromosome, so R133C, R168X and R270X alleles are homozygous, mimicking the 50 percent mutant cell fraction in female patients. Reprogramming of female patient fibroblasts into iPSCs induces erosion of X-chromosome inactivation (XCI), resulting in heterozygous *MECP2* expression.^42,43^ Quantification of XIST transcript levels confirmed XCI loss in all eight iPSC lines (Figure S5). Quantitative RT-PCR demonstrated that *ZFP42* overexpression is specific to homozygous truncating alleles, as heterozygous iPSCs retained WT-level *ZFP42*, similar to *POU5F1* (Figure 3D). To investigate whether Rett hESC acquire a naïve-like transcriptional program, we interrogated the top 20 naïve-enriched markers (Figure 3E).^38^ Eighteen of these markers were elevated in mutant lines, including key markers like *PRDM14* and *SALL4* in R168X and R270X, whereas R133C selectively increased a different subset of naïve-associated genes despite exhibiting baseline *ZFP42* levels (Figure 3E). To assess whether *MECP2* loss accelerates proliferation, a hallmark of naïve state,^44,45^ we performed cell-cycle analysis by assigning single-cell transcriptomes to G₁, S or G₂/M phases (Figure 3F-G). WT hESCs spend ∼82 percent of time in S+G₂/M, whereas R168X and R270X maintain or expand this replicative pool (81 percent and 86 percent) and R133C shows an intermediate profile (71 percent) (Figure 3G). These results indicate that truncating *MECP2* mutations shorten G₁ and extend S+G₂/M occupancy, driving a hyperproliferative, naïve-like cell cycle profile. To directly quantify growth kinetics, we captured live hESC colony growth. *MECP2* mutants exhibited a pronounced proliferation advantage relative to WT (Figure 3H). Taken together, these data indicate that MECP2 loss shifts hESCs toward a naïve-like, hyperproliferative state and highlight ZFP42 as a critical outlier for mechanistic investigation.

### Convergent *EMX1* repression in *MECP2*-mutant stem cells drives cortical lineage imbalance

To pinpoint early, mutation-shared transcriptional defects that could prefigure downstream developmental abnormalities, we first performed bulk differential expression analysis on undifferentiated hESCs carrying the R133C, R168X or R270X *MECP2* alleles. This comparison yielded two compact, convergent gene sets with 143 transcripts down-regulated and 139 up-regulated in all mutants relative to WT (Figure 4A). *EMX1*, a LIM-homeodomain transcription factor that specifies dorsal telencephalic progenitors, emerged among the most strongly repressed genes (Figure 4A).^46,47^ Given its essential role in cortical fate commitment, we queried whether *EMX1* repression generalizes across patient material. Quantitative RT-PCR in two independent control iPSC lines and eight female Rett iPSC lines confirmed a consistent loss of *EMX1* expression in patient iPSCs (Figure 4B), indicating that diminished *EMX1* is neither allele-specific nor confined to male hESCs. To establish the developmental context of EMX1 action, we mined reference single-cell atlases spanning mouse embryonic cortex (E12–E15), human fetal cortex (8– 20 weeks), and human brain organoids (10 weeks).^48^ In all three systems, *EMX1* transcripts are restricted to apical radial glia (aRG) and nascent excitatory neurons, positioning EMX1 at the apex of cortical excitatory lineage specification (Figure 4C). This observation prompted us to ask whether early EMX1 loss translates into altered lineage allocation in RTT at later stages. We therefore generated 3-months old unguided cerebral organoids from each genotype and profiled them using single-nucleus RNA sequencing, an age when both excitatory and inhibitory neurons, as well as glial lineages, are readily detectable. UMAP projection separated nuclei into six canonical populations: outer radial glia (oRG), intermediate progenitor cells (IPC), oligodendrocyte lineage, excitatory neurons, inhibitory neurons and astrocytes (Figure 4D). Cell-type quantification revealed pronounced mutation-specific shifts in lineage allocation compared to WT (Figure 4E). Control organoids were dominated by oligodendroglial derivatives (∼40–50%) with modest inhibitory and minimal excitatory neuron fractions. In contrast, R133C organoids exhibited an expanded inhibitory compartment (∼50–55 %), R168X organoids accumulated oligodendrocyte progenitors (30–35 %) while retaining few excitatory neurons (<5 %), and R270X organoids showed a striking predominance of excitatory neurons (70–80 %) with minimal glial contribution. These data indicate that *MECP2* mutations perturb cortical cell-type balance in distinct, allele-specific patterns (Figure 4E). Together, these findings delineate a coherent trajectory linking early, *MECP2*-dependent repression of *EMX1* with later biases in cortical lineage outcome.

**Figure 4.**
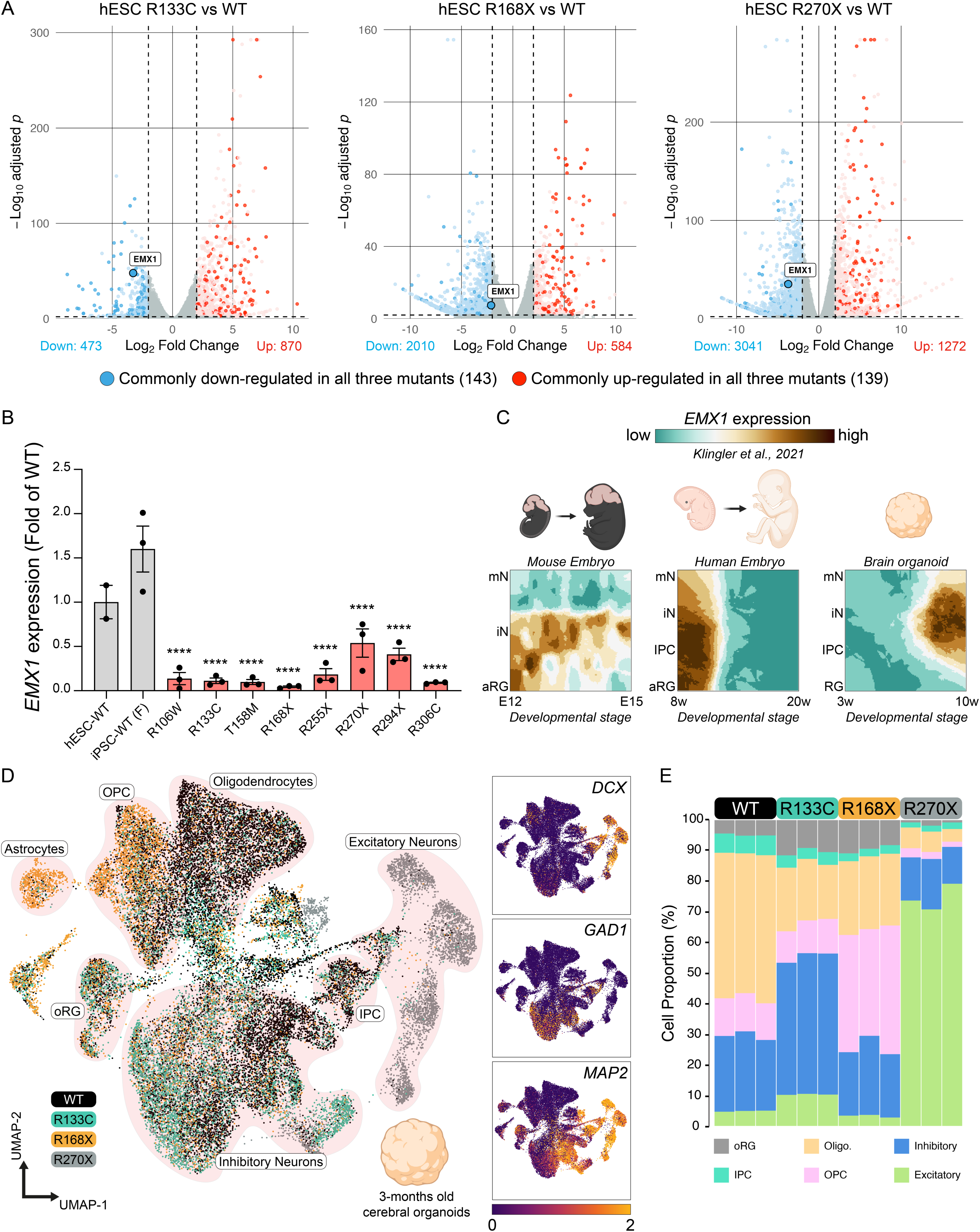
Convergent *EMX1* repression links early defects to reduced excitatory-neuron output in Rett organoids. **A.** Volcano plots of differential expression (log₂ fold-change vs –log₁₀ adjusted p) for each mutant hESC line relative to WT. Genes down-regulated in all three mutants are highlighted in blue (n = 143) and up-regulated genes in red (n = 139). The cortical progenitor determinant *EMX1* is indicated in each plot. Computed from bulk RNA sequencing matrix. **B.** Quantitative RT-PCR validation of *EMX1* expression across pluripotent lines: WT hESCs, control iPSC line (grey), and eight Rett iPSC lines bearing the indicated *MECP2* alleles (red). Bars represent mean ± s.e.m.; asterisks denote one-way ANOVA with Dunnett’s post-hoc test versus WT hESC (**** *P* < 0.0001). F: Female. **C.** Reference single-cell atlases showing spatial localization of *EMX1* transcripts (teal–brown scale) in mouse embryonic cortex (E12–E15), human fetal cortex (8–20 weeks) and 10-week brain organoids. aRG, apical radial glia; IPC, intermediate progenitor cell; iN, immature neuron; mN, mature neuron. **D.** Left, UMAP embedding of 25,132 single nuclei from three-month cerebral organoids (N=3 per genotype), colored by genotype and annotated into six major populations: outer radial glia (oRG), intermediate progenitor cells (IPC), excitatory neurons, inhibitory neurons, oligodendrocyte, oligodendrocyte progenitor cells (OPC) and astrocytes. Right, feature plots of lineage markers *DCX* (excitatory), *GAD1* (inhibitory) and *MAP2* (pan-neuronal). **E.** Stacked bar chart displaying the proportional composition of each cell type per organoid across WT and the three mutant genotypes.

## Discussion

By integrating stage-resolved bulk and single-cell transcriptomics with functional assays in isogenic human ESCs, RTT patient-derived iPSCs and long-term cerebral organoids, we identify a shared, MECP2-dependent transcriptional program that is already perturbed in the hypomethylated pluripotent state. Three convergent features emerge. First, a discrete set of 282 transcripts is consistently mis-regulated across three representative *MECP2* mutations, indicating that MECP2 influences a restricted yet coherent gene network even before neural induction. Second, truncating alleles (R168X, R270X) elicit a naïve-like molecular signature, marked by *ZFP42*, *PRDM14* and *SALL4* elevation, and a concomitant shortening of G1 that confers a proliferation advantage. Third, all mutants display strong repression of the dorsal telencephalic determinant *EMX1*, and this early lesion is followed by a quantitative deficit in excitatory-neuron production in three-month cerebral organoids. Collectively, our data situate the origin of RTT-relevant defects at the pluripotent stage and outline a trajectory through which early transcriptional noise may culminate in cortical-circuit imbalance.

### Early transcriptional disruption and its developmental implications

The detection of genotype-specific variance on principal component 2 as early as the ESC stage (Figure 1B) argues that MECP2 regulates transcription independently of the canonical methyl-DNA pathway, consistent with recent reports of methylation-independent MECP2 binding to enhancer-like hotspots.^16,17^ The convergent mis-regulation of *S100A6* and *SLC17A7* across all stages (Figure 1C) further suggests that cytoskeletal organization and glutamatergic signaling are primed for later dysfunction before lineage commitment occurs. While our machine-learning classifiers could readily decode developmental stage, they also highlighted an anomalous “early surge–late collapse” pattern for neuronal genes in mutants (Figure 2), hinting at mistimed engagement of neurogenic program. Such temporal mis-alignment may help reconcile seemingly contradictory findings in mouse models where *Mecp2* loss produces both premature and delayed neuronal maturation.^49,50^ Future time-resolved chromatin profiling, ideally combined with base-resolution methylome maps, will be needed to decipher whether the observed transcriptomic shifts reflect direct MECP2 occupancy or secondary chromatin remodeling events.

### A putative shift toward naïve pluripotency

Up-regulation of *ZFP42*, *PRDM14*, *TACSTD2* and other naïve markers, together with a shortened G1, points to a partial drift from the primed toward the naïve state. Naïve human PSCs are characterized by reduced H3K27me3 deposition, global DNA hypomethylation, biallelic expression of X-linked genes and elevated oxidative phosphorylation.^38^ Our data fulfil two of these criteria (*ZFP42*/*PRDM14* induction and G1 compression) yet do not show broad deregulation of canonical pluripotency factors or detectable copy number variation (Figure 3 and S3). Thus, the shift appears partial and may reflect a “metastable” intermediate rather than a full reset to an embryonic-like ground state. To refine this interpretation, assays that probe functional hallmarks of naïve cells, such as competence for trophectoderm induction, transposon activation profiles, global hydroxymethylcytosine levels or enhancer re-wiring, should be performed. Equally important will be to determine whether the naïve-like drift is cell-autonomous or emerges from subtle culture-selection advantages conferred by hyper-proliferation.

### EMX1 as an early bottleneck for cortical excitatory fate

Among the 143 convergently down-regulated genes, *EMX1* stands out because of its restricted expression in apical radial glia and nascent excitatory neurons, two populations that seed the cortical excitatory lineage. Our single-nucleus RNA sequencing data reveal that *EMX1* repression is followed by a reduced fraction of excitatory neurons and a compensatory expansion of inhibitory or glial lineages in three-month organoids. Although correlative, this trajectory is consistent with mouse studies showing that Emx1 knock-out skews cortical progenitors toward ventral or glial fates.^51^ Elucidating causality will require gain-of-function experiments in *MECP2*-mutant ESCs and organoids. Rescue of excitatory-neuron output would provide compelling evidence that EMX1 is a pivotal effector of early MECP2 function. Conversely, single-cell multi-omics combining chromatin accessibility and transcript profiling could clarify whether *EMX1* repression is direct (via MECP2 occupancy) or indirect via intermediate transcription factors.

### Translational outlook

On the therapeutic front, the partial naïve-like drift and early *EMX1* repression raise the possibility that interventions targeting chromatin state or lineage-bias pathways might complement post-natal gene-replacement strategies. Epigenetic drugs that stabilize the primed state or small molecules that up-regulate *EMX1* could, in principle, normalize lineage allocation if applied at appropriate developmental windows. Whether such interventions are feasible, safe and effective *in vivo* remains an open question.

## Limitations of the Study

The concept that neurodevelopmental disorders originate from perturbations in pluripotent or early progenitor states is gaining traction in Fragile X, 22q11.2 deletion syndrome and autism spectrum disorder models.^52–54^ Our findings position RTT within this framework and suggest that therapeutic windows may extend into very early developmental stages. However, translating these insights to patients will require caution. First, our study relies on *in vitro* models that, while genetically controlled, may not fully capture *in vivo* morphogen gradients or cell-extrinsic cues. Second, the male ESC system, chosen to avoid mosaic X-inactivation, may over-represent homozygous effects that are present in only ∼50 % of cells in female patients. Employing isogenic female ESCs with inducible *XIST* or using allele-specific reporters could address this limitation.

## Acknowledgments

We thank Dr. Rudolf Jaenisch and his team for sharing key reagents and providing strategic guidance. We thank Drs. Graziella Di Cristo, Elsa Rossignol, and Serge McGraw for their insightful guidance; Dr. Eric Samarut for critical review of the manuscript; all members of the Flamier laboratory for their contributions; Basma Benabdallah and the CHU Sainte-Justine iPSC core facility; and Nicholas Geoffrion and the bioinformatics core facility for essential technical support. This work was funded by the Canada Brain Research Fund (CBRF)—a partnership between Health Canada and the Brain Canada Foundation (Future Leaders Program); the Canadian Institutes of Health Research; the Canadian Stem Cell Network Jump Start ECR Program; the CHU Sainte-Justine Foundation; the Fonds de Recherche du Québec–Santé (FRQS); and the Rett Syndrome Research Trust (for providing Rett iPSC lines). Additional support was provided by the Partenariat en épilepsie entre l’Institut Imagine de Paris et le CHU Sainte-Justine. We also thank the International Rett Syndrome Foundation (IRSF) Research Independence Award (to Y.L.).

## Author Contributions

A.F. conceived the project, obtained funding, and coordinated all research activities (Conceptualization, Funding acquisition, Project administration). M.G, M.B., J.G., L.L, T.R, E.G., M.B. and A.F. generated and curated the datasets, carried out the majority of wet-lab experiments, performed data validation, and prepared figures (Investigation, Data curation, Validation, Visualization). Y.L. and M.M. supplied key reagents, maintained cell lines, and contributed to experimental execution (Resources, Investigation). L.L., L.A., and A.F. supervised the work and provided strategic guidance (Supervision). A.F. wrote the initial manuscript draft, and all authors revised and edited the final version (Writing – original draft; Writing – review & editing). All authors read and approved the manuscript.

## Declaration of Generative AI and AI-assisted Technologies in the Writing Process

During the preparation of this work, the authors used OpenAI ChatGPT Large Language Model o3 to enhance the clarity of the text. The authors reviewed and edited the content and take full responsibility.

