## Supplemental Information for "*MECP2* Mutations Rewire Human ESC Fate and Bias Cortical Lineage Commitment"

### SUPPLEMENTARY INFORMATION

Number of supplementary figures: 5

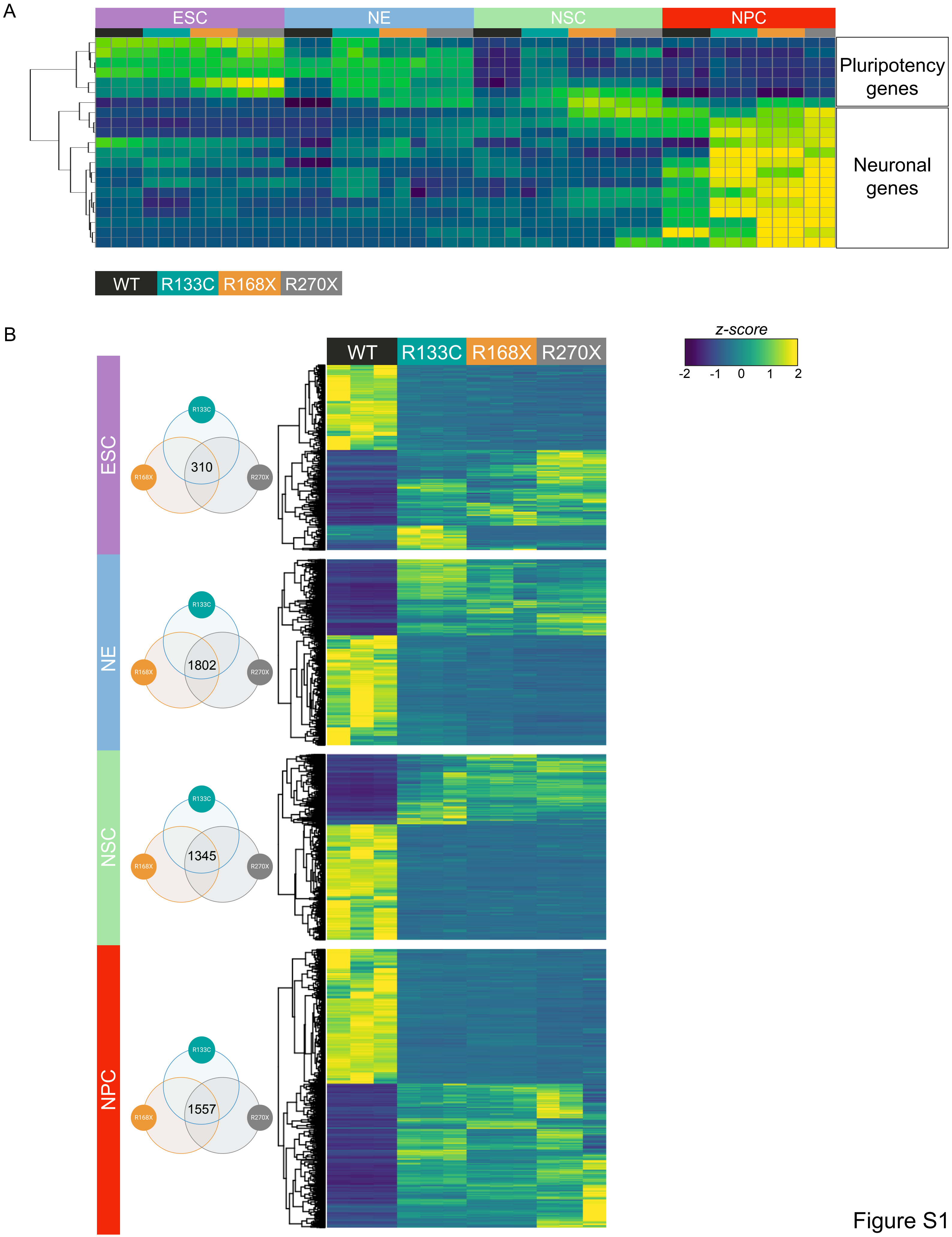
A
ESC
NE
NSC
NPC
Pluripotency genes
Neuronal genes
WT
R133C
R168X
R270X
B
z-score
-2
-1
0
1
2
ESC
R133C
R168X
R270X
310
NE
1802
NSC
1345
NPC
1557

Figure S1

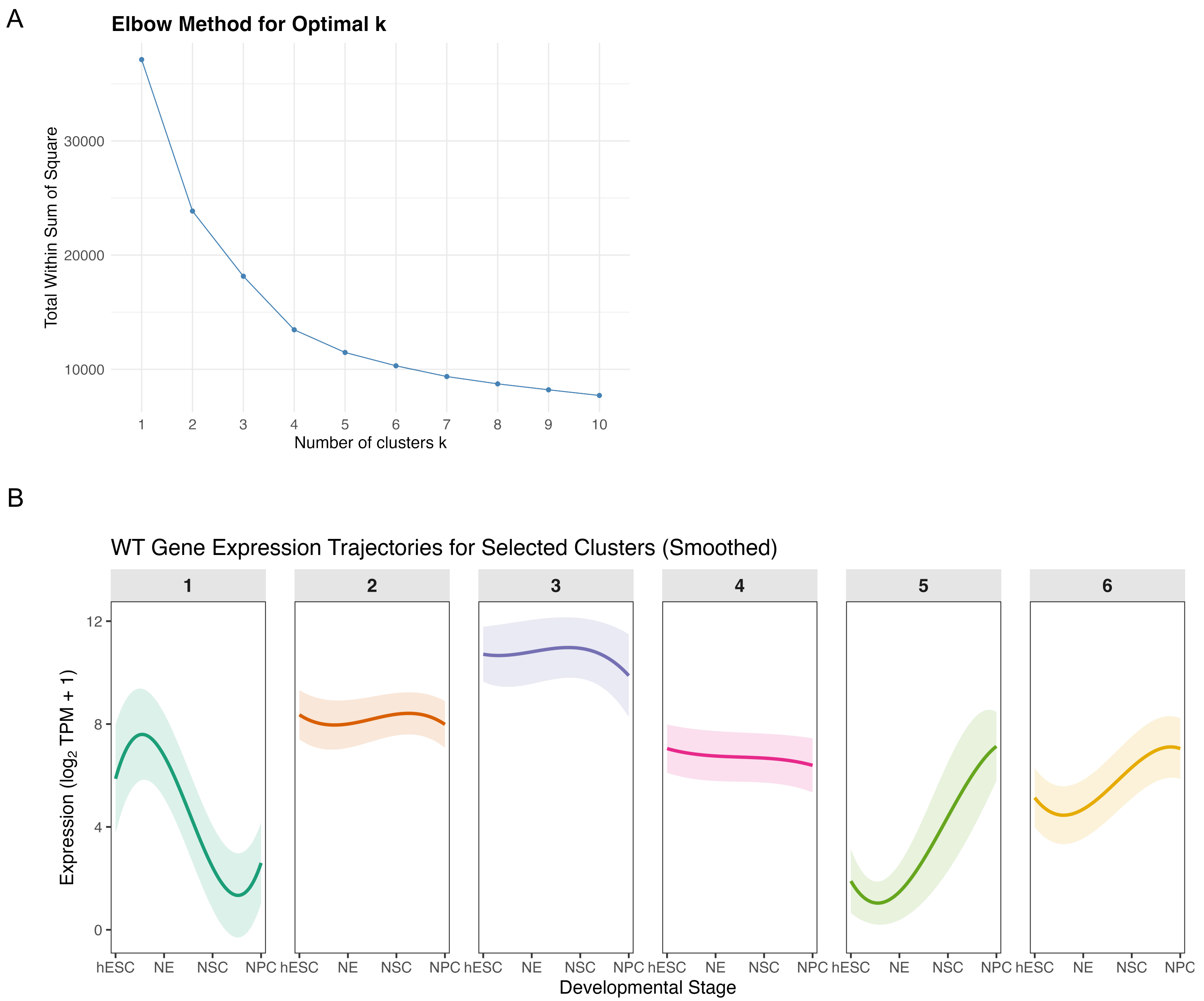
A
Elbow Method for Optimal k
Total Within Sum of Square
30000
20000
10000
1
2
3
4
5
6
7
8
9
10
Number of clusters k
B
WT Gene Expression Trajectories for Selected Clusters (Smoothed)
1
2
3
4
5
6
Expression ($log_2$ TPM + 1)
12
8
4
0
hESC
NE
NSC
NPC
Developmental Stage

Figure S2

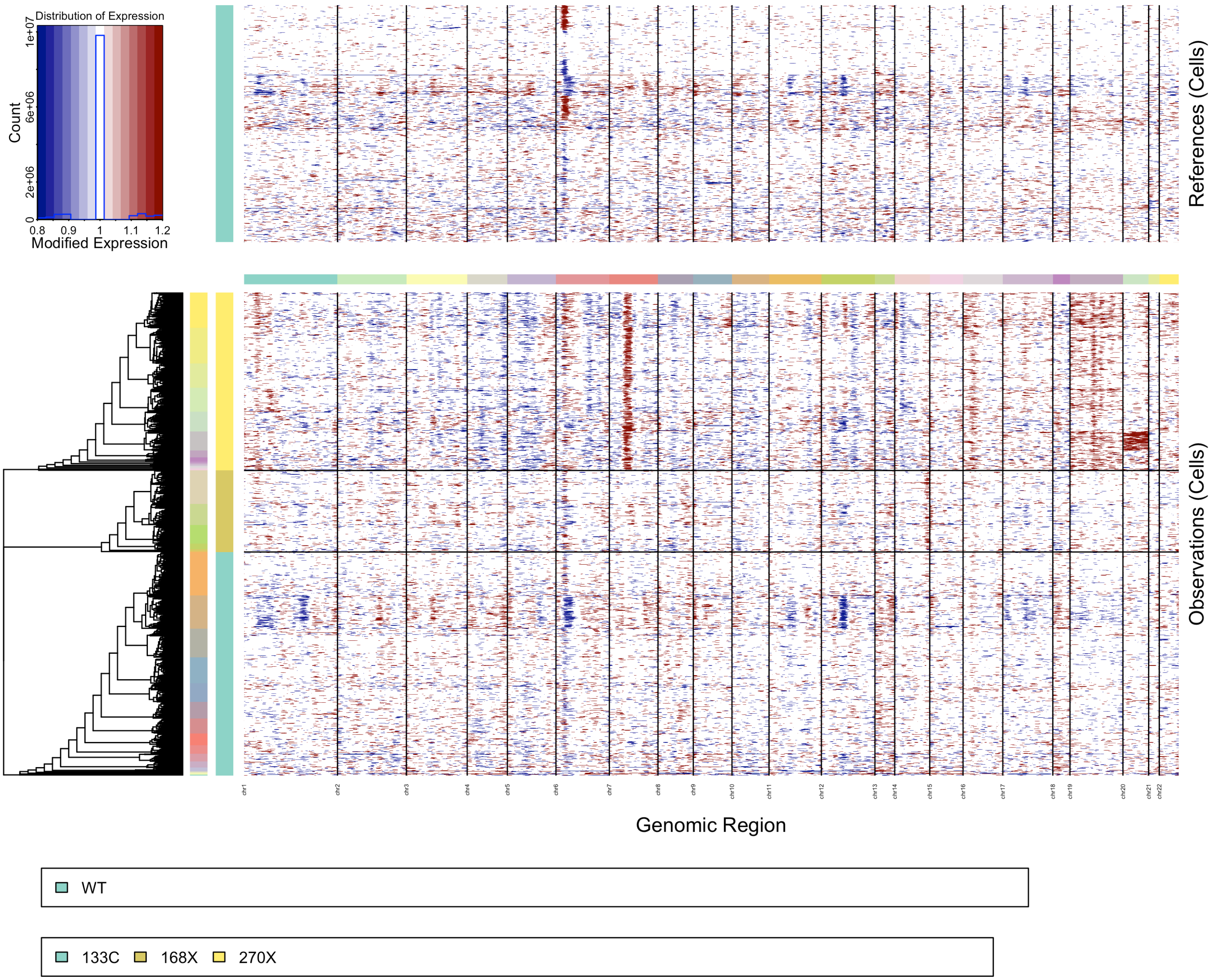
Distribution of Expression
Count
Modified Expression
0.8 0.9 1 1.1 1.2
0 2e+06 6e+06 1e+07
References (Cells)
Observations (Cells)
chr1 chr2 chr3 chr4 chr5 chr6 chr7 chr8 chr9 chr10 chr11 chr12 chr13 chr14 chr15 chr16 chr17 chr18 chr19 chr20 chr21 chr22
Genomic Region
WT
133C 168X 270X

Figure S3

A

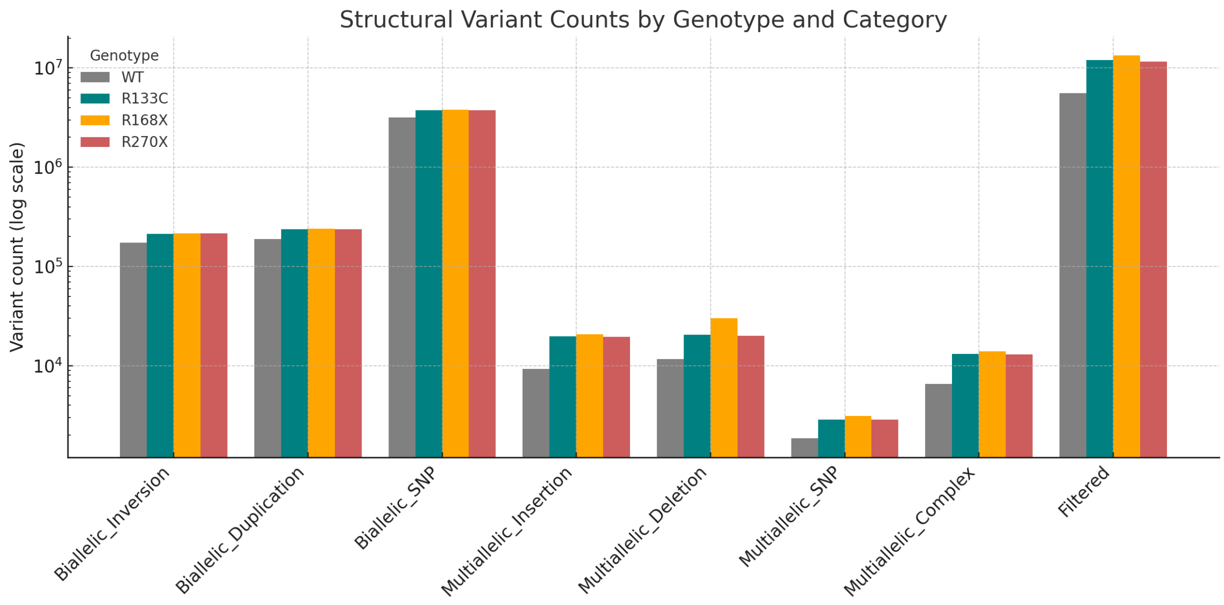
Structural Variant Counts by Genotype and Category
Genotype
WT
R133C
R168X
R270X
Variant count (log scale)
$10^7$
$10^6$
$10^5$
$10^4$
Biallelic_Inversion
Biallelic_Duplication
Biallelic_SNP
Multiallelic_Insertion
Multiallelic_Deletion
Multiallelic_SNP
Multiallelic_Complex
Filtered

B

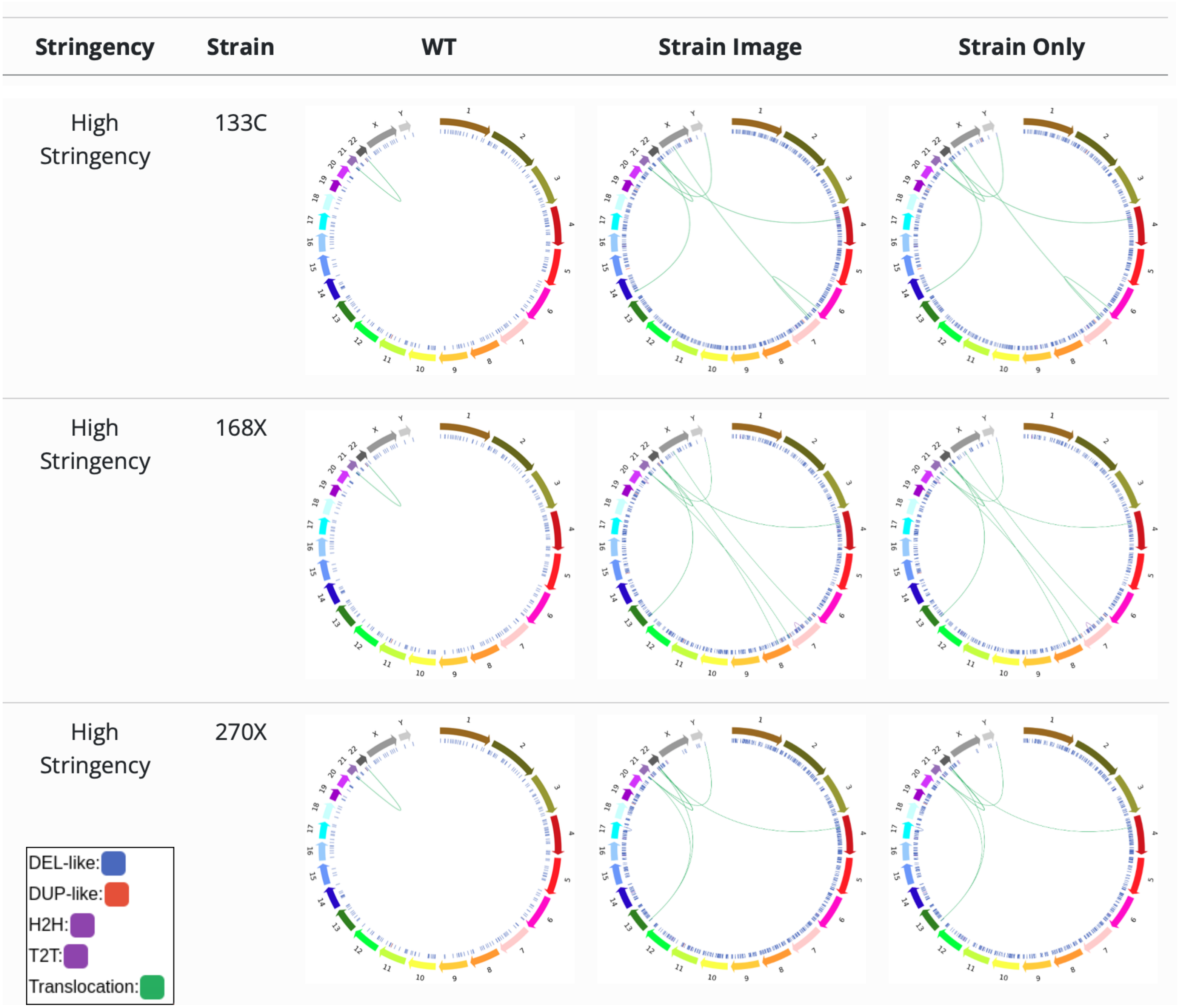
Stringency
Strain
WT
Strain Image
Strain Only
High Stringency
133C
High Stringency
168X
High Stringency
270X
DEL-like:
DUP-like:
H2H:
T2T:
Translocation:

Figure S4

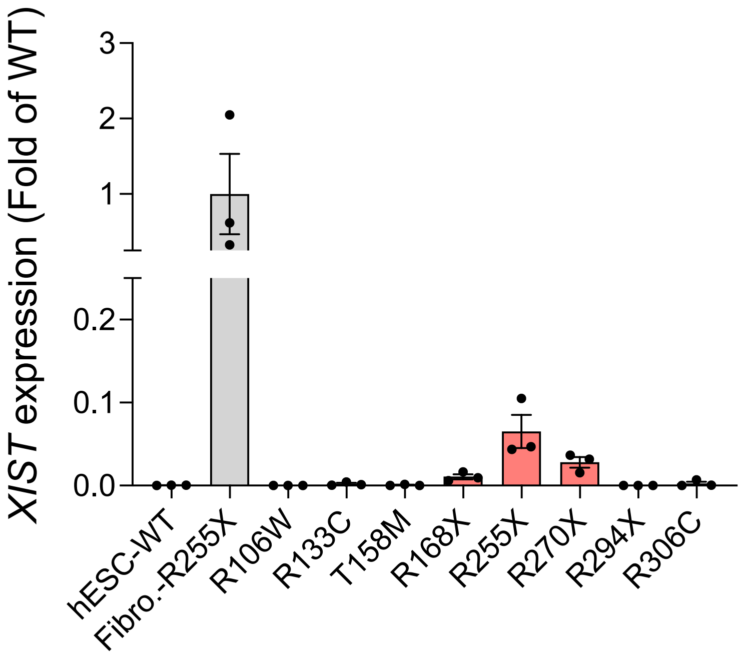

XIST expression (Fold of WT)
3
2
1
0.2
0.1
0.0
hESC-WT
Fibro.-R255X
R106W
R133C
T158M
R168X
R255X
R270X
R294X
R306C


Figure S5

### Figure Legends

**Figure S1.** Related to Figure 1

**A.** Z-score heat-map of curated pluripotency (top block) and neuronal (bottom block) marker genes across the four differentiation stages (ESC, NE, NSC, NPC; color bar above) for WT and the three mutant hESC lines (key below). Replicate columns are ordered by hierarchical clustering (Ward's method, Euclidean distance).

**B.** For each stage, Venn diagrams (left) enumerate significantly dysregulated genes common to all mutants (Wald test, DESeq2; $\log_2$FC $> 2$, padj $< 0.01$). The corresponding union set is visualized as a heat-map (right; same scaling as in A) with rows clustered by Pearson correlation.

**Figure S2.** Related to Figure 2

**A.** Elbow plot of total within-cluster sum of squares across $k = 1–10$; $k = 6$ was chosen as the inflection point for all further k-means analyses.

**B.** Smoothed (loess) expression trajectories of WT genes belonging to the six selected clusters (shaded 95 % confidence band; y-axis, $\log_2$ TPM + 1; x-axis, developmental stage).

**Figure S3.** Related to Figure 3

Top panel, reference WT cells; bottom panel, combined R133C, R168X and R270X observations. Each column represents a genomic region ordered chromosomally from 1 to Y (black tick marks) and each row a single cell, clustered by Euclidean distance. Red denotes inferred gains, blue denotes losses (scale bar, left). Color strips mark genotype identity of observation cells. Dendrogram shows hierarchical relationships among mutant cells.

**Figure S4.** Related to Figure 3

**A.** Grouped bar chart of structural-variant (SV) counts binned by type ($\log_{10}$ scale) for WT and mutants at high-stringency Sniffles2 settings (≥8 supporting reads, ≥150 bp).

**B.** Circos plots of SVs retained after comparison with the WT call-set ("Strain Only") for each mutant. Outer track, chromosomes 1–22, X, Y; inner chords, high-confidence SVs colored by class (legend, bottom left). Central column shows WT reference, confirming removal of shared variants.

**Figure S5.** Related to Figure 3

Relative *XIST* expression measured by RT-qPCR (normalized to fibroblasts) in control hESCs (grey), primary fibroblasts from patient R255X (left, grey bar), and eight Rett patient-derived iPSC lines (red bars). Bars, mean of three biological replicates; dots, individual values; error bars, ± s.e.m. Loss of *XIST* confirms erosion of X-chromosome inactivation in patient iPSCs.

### KEY RESOURCES TABLES

| REAGENT OR RESOURCE | SOURCE | IDENTIFIER |
|---|---|---|
| **Chemicals, peptides, and recombinant proteins** | | |
| Corning Matrigel Basement Membrane Matrix, LDEV-free | Corning | 354234 |
| Corning Matrigel Growth Factor Reduced (GFR) Basement Membrane Matrix, Phenol Red-free, LDEV-free | Corning | 356231 |
| Dimethyl sulfoxide | Sigma-Aldrich | D8418 |
| DMEM/F12 | Wisent Inc | 319-090 CL |
| Gentle Cell Dissociation Reagent | Stem Cell Technologies | 100-0485 |
| KnockOut Serum Replacement | Gibco | 10828010 |
| mTeSR Plus | Stem Cell Technologies | 100-0276 |
| mTeSR Plus Supplement | Stem Cell Technologies | 100-0275 |
| Pen Strep Glutamine (100X) | Gibco | 10378-016 |
| Plasmocin Prophylactic | InvivoGen | ant-mpp |
| PowerUp SYBR Green Master Mix for qPCR | Applied Biosystems | A25742 |
| QIAshredder | Qiagen | 79656 |

| | | |
|---|---|---|
| qScript cDNA SuperMix | Quantabio | 95048-100 |
| ReleSR | Stem Cell Technologies | 100-0483 |
| STEMDiff SMADI Neural Induction Supplement | Stem Cell Technologies | 08580 |
| STEMDiff Neural Induction | Stem Cell Technologies | 05835 |
| Trypan blue Trypan blue | Thermo Fisher | T10282 |
| UltraPure DNase/RNase-Free Distilled Water | Invitrogen™ | 10977-015 |
| Y-27632 (Dihydrochloride) | Stem Cell Technologies | 72304 |
| **Critical commercial assays** | | |
| DNA 1000 Kit | Agilent Technologies | 5067-1504 |
| EVERCODE Cell Fixation v2 | Parse Bioscience Inc | ECF2101 |
| EVERCODE Nuclei Fixation v2 | Parse Bioscience Inc | ECF2103 |
| EVERCODE WT Mini v2 | Parse Bioscience Inc | ECW02115 |
| EVERCODE WT v2 | Parse Bioscience Inc | ECW02135 |
| NEBNext High Input Poly(A) mRNA Isolation Module | New England Bioloabs | E3370S |
| NEBNext Ultra II Directional RNA Library Prep Kit for Illumina | New England Bioloabs | E7760L |
| PureLink RNA Mini Kit | Thermo Fisher Scientific | 12183018A |
| QuantiFluor dsDNA system | Promega Corporation | E2670 |
| RNeasy Mini Kit | Qiagen | 74106 |
| STEMDiff Cerebral Organoid Kit | Stem Cell Technologies | 08570 |
| **Deposited data** | | |
| RNA sequencing | This study | GEO: GSE303838 |
| Single-nucleus RNA sequencing | This study | GEO: GSE303977 |
| Single-cell RNA sequencing | This study | GEO: GSE303813 |
| Rett Data Explorer | This study | https://e37e9c0c0387.ngrok-free.app |
| **Experimental models: Cell lines** | | |
| hESC-WIBR1 R133C MECP2-GFP | Liu, Y. *et al.* [1] | N/A |
| hESC-WIBR1 R168X MECP2-GFP | Liu, Y. *et al.* [1] | N/A |
| hESC-WIBR1 R270X MECP2-GFP | Liu, Y. *et al.* [1] | N/A |
| hESC-WIBR1 WT MECP2-GFP | Liu, Y. *et al.* [1] | N/A |
| hiPSC 7095 | This study | N/A |
| hiPSC Col7A1 WT female | This study | N/A |
| hiPSC p.R106W | RSRT iPSC Collection Coriell | N/A |
| hiPSC p.R133C | RSRT iPSC Collection Coriell | N/A |
| hiPSC p.T158M | RSRT iPSC Collection Coriell | N/A |
| hiPSC p.R168X | RSRT iPSC Collection Coriell | N/A |
| hiPSC p.R255X | RSRT iPSC Collection Coriell | N/A |

| | | |
|---|---|---|
| hiPSC p.R270X | RSRT iPSC Collection Coriell | N/A |
| hiPSC p.R295X | RSRT iPSC Collection Coriell | N/A |
| hiPSC p.R306C | RSRT iPSC Collection Coriell | N/A |
| **Software and algorithms** | | |
| DESeq2 | Love *et al.* [2] | https://bioconductor.org/packages/release/bioc/html/DESeq2.html<br>RRID:SCR_015687 |
| GraphPad Prism GraphPad Software V.10 | N/A | https://www.graphpad.com/scientific-software/prism/<br>RRID:SCR_002798 |
| Fiji image processing package.76 | Schindelin *et al.* [3] | http://fiji.sc<br>RRID:SCR_002285 |
| featureCounts | Liao et al. [4] | https://subread.sourceforge.net/featureCounts.html<br>RRID:SCR_012919 |
| STAR | Dobin et al. [5] | https://hbctraining.github.io/Intro-to-rnaseq-hpc-O2/lessons/03_alignment.html<br>(RRID:SCR_004463) |
| FASTQC | N/A | https://www.bioinformatics.babraham.ac.uk/projects/fastqc/<br>(RRID:SCR_014583) |
| SAMtools | Li et al. [6] | https://www.htslib.org/<br>RRID:SCR_002105 |
| Seurat V.5 | Hao et al. [7] | https://satijalab.org/seurat/<br>(RRID:SCR_016341) |
| Trim Galore! | N/A | https://github.com/FelixKrueger/TrimGalore<br>RRID:SCR_011847 |
| clusterProfiler | Yu et al., [8] | https://bioconductor.org/packages/devel/bioc/html/clusterProfiler.html<br>(RRID:SCR_016884) |

| | | |
|---|---|---|
| RandomForest Package in R | Breiman [9] | https://cran.r-project.org/web/packages/randomForest/index.html (RRID:SCR_015718) |
| Keras | N/A | https://github.com/rstudio/keras (RRID:SCR_026159) |
| TensorFlow | N/A | https://cran.r-project.org/web/packages/tensorflow/index.html (RRID:SCR_016345) |

### RESOURCE AVAILABILITY

*Lead Contact*

*Materials Availability*

All unique, stable reagents generated in this study are available from the lead contact upon reasonable request, contingent on completion of a Materials Transfer Agreement.

*Data and Code Availability*

Genomic data are available through public repository (bulk RNAseq: GSE303838; scRNAseq: GSE303813; snRNAseq: GSE303977). A data explorer app is available at: https://e37e9c0c0387.ngrok-free.app

All other raw data and codes are available upon request to the lead contact.

### METHOD DETAILS

#### Pluripotent Stem Cell Culture

Human embryonic stem cells (hESCs) WIBR1 were maintained under feeder-free conditions. Three CRISPR/Cas9-edited male hESC lines carrying the recurrent Rett-syndrome mutations R133C, R168X, and R270X were used alongside the isogenic wild-type (WT) parental line as previously reported[1]. In addition, eight female patient-derived induced pluripotent stem-cell (iPSC) lines harbouring distinct *MECP2* variants (R106W, R133C, T158M, R168X, R255X, R270X, R294X, R306C) were obtained from the RSRT iPSC Collection (Coriell Institute). For routine culture, cells were thawed onto tissue-culture plates coated with hESC-qualified Matrigel (Corning, #354277) and maintained in mTeSR Plus medium (Basal Medium #100-0274 supplemented 1:5 with mTeSR Plus 5× Supplement, #100-2075; STEMCELL Technologies). Cultures were incubated at 37 °C in 5 % $CO_2$ and medium was refreshed every 24 h. Cells were passaged every 3–4 days using ReLeSR (STEMCELL Technologies, #100-0483); 10 μM Y-27632 ROCK inhibitor (STEMCELL Technologies, #72308) was added for 24h after passaging to enhance survival.

#### Neuronal differentiation

hESCs were plated on three well of a 6-well plate coated with Growth Factor Reduced Matrigel using ReLeSR (STEMCELL Technologies, #100-0483). After 24h (around 70% confluence) cells were incubated with Neural induction media with SMAD inhibitors (STEMCELL Technologies, #08580) with daily full media change. Differentiated cells were stopped 3 days (Neuroectodermal stage), 7 days (Neural stem cell stage) and 21 days (Neural progenitor cell stage) post induction.

#### Cell-growth assay

Individual hESC lines were seeded onto 6-well plates pre-coated with hESC-qualified Matrigel (Corning, #354277). After a 24-h attachment period, plates were transferred to an Incucyte S3 live-cell imager and kept under standard culture conditions (37 °C, 5 % $CO_2$). Nine non-overlapping phase-contrast images per well were captured every 4 h for 72 h. For each well three independent colonies of comparable initial size across genotypes were manually selected, and its perimeter was measured at every time point in ImageJ. Growth curves were generated by plotting the perimeter of the same colony over the 72-h imaging window.

#### Unguided cerebral-organoid differentiation

Unguided cortical organoids were generated from WT, R133C, R168X and R270X hESCs with the STEMdiff Cerebral Organoid Kit (STEMCELL Technologies, #08570). Briefly, hESCs were dissociated to single cells with ReLeSR, counted, and seeded at $9 \times 10^3$ cells per well in low-attachment, V-bottom 96-well plates supplied with the kit (day 0). Aggregates were cultured in Embryoid Body Medium for 5 days, with half-medium changes on days 2 and 4, to promote uniform spheroid formation. On day 5, spheroids were transferred to neural induction medium in ultra-low-attachment 24-well plates (three spheroids per well). After 48 h (day 7), each spheroid was embedded in a 30-µl dome of growth-factor-reduced Matrigel and moved to maturation medium in a Celtron orbital shaker (INFORS HT, #I69222) operating at 75 rpm inside a humidified incubator (37 °C, 5 % $CO_2$). Medium was exchanged every 3–4 days for the duration of the culture. To limit central hypoxia and necrosis, organoids were sectioned under sterile conditions beginning at day 60. Using a Leica VT1000 S vibratome fitted with a sterile blade, each organoid was bisected or trisected (300–400 µm slices) every 2–3 weeks and individual fragments were returned to the shaker in fresh maturation medium. Cultures were harvested at 4 months for downstream single-nucleus RNA-seq and histological analyses.

**Reverse-Transcription quantitative PCR (RT-qPCR)**

Total RNA was isolated from each hiPSC line in triplicate (n = 3 biological replicates per genotype) with the RNeasy Mini Kit (Qiagen, #74106) on a QIAcube automated workstation. For cDNA synthesis, 500 ng of RNA were reverse-transcribed using qScript cDNA SuperMix (Quantabio, #95048-100) on a SimpliAmp thermal cycler, following the manufacturer's protocol. The resulting cDNA was diluted 1:8 with nuclease-free water and used as template for SYBR Green qPCR. Reactions (10 µL total) were assembled in 96-well plates as follows: 5 µL PowerUp SYBR Green Master Mix (Thermo Fisher Scientific), 0.5 µL forward primer (10 µM), 0.5 µL reverse primer (10 µM), 3 µL UltraPure DNase/RNase-free distilled water, and 1 µL diluted cDNA.

Primer sequences were:

EMX1_F: ACCGGGACCCTCTCCATTT

EMX1_R: GCTTCTGCCGTTTGTACTTTGT

ZFP42_F: AGAAACGGGCAAAGACAAGAC

ZFP42_R: GCTGACAGGTTCTATTTCCGC

XIST_F: CCAGGCAATCTGCTCTGGAA

XIST_R: ATGCTGACTACCCAAAGCCC

POU5F1_F: CCCCAGGGCCCCATTTTGGTACC

POU5F1_R: ACCTCAGTTTGAATGCATGGGAGAGC

RPLP0_F: AGCCCAGAACACTGGTCTC

RPLP0_R: ACTCAGGATTTCAATGGTGCC

Thermal cycling conditions followed the Master Mix guidelines (two-step protocol with melt-curve analysis). All samples were run in technical triplicate, and relative expression was calculated by the ΔΔCt method after normalization to the geometric mean of *RPLP0*.

**Bulk RNA Sequencing**

Total RNA was extracted in biological triplicate (n = 3 per line) with the PureLink RNA Mini Kit (Invitrogen, #12183018A) at four matched time points: day 0, day 3, day 7 and day 21 for WT and the three point-mutant lines. Polyadenylated RNA was purified from each triplicate with the NEBNext Poly(A) mRNA Magnetic Isolation Module (New England Biolabs, #E3370S). One microlitre of the eluate was quantified on a NanoDrop spectrophotometer. In total, 48 barcoded libraries were prepared with the NEBNext Ultra II Directional RNA Library Prep Kit for Illumina (NEB, #E7760L) according to the manufacturer's instructions. Libraries were sequenced on Illumina NovaSeq X Plus instruments using paired-end 50-bp reads (PE50) with at least 40 million reads per sample.

**Bulk RNA Sequencing analysis**

Raw paired-end FASTQ files were quality-checked with FastQC (v0.11.9) and summarised with MultiQC. Adapter sequences and bases with Phred < 20 were removed using Trimmomatic (v0.39; settings ILLUMINACLIP:2:30:10, SLIDINGWINDOW:4:20, MINLEN:36). Cleaned reads were aligned to the GRCh38/hg38 reference (Ensembl 109 annotation) with STAR (v2.7.11a) in two-pass mode (--twopassMode Basic) and sorted BAMs were produced (--outSAMtype BAM SortedByCoordinate). Gene-level counts were obtained with featureCounts (v2.0.2; options -p -B -C --primary -s 0) and imported into R (4.3.2). Low-abundance features (row-sums ≤ 1) were discarded, and a DESeq2 (v1.40.2) object was created with the design formula ~ Genotype + Time. After dispersion estimation and Wald testing, contrasts were extracted for each mutant versus WT at each time point; genes with $\log_2 FC > 2$ and Benjamini–Hochberg–adjusted $p < 0.01$ were considered differentially expressed. Variance-stabilised counts (vst, blind = FALSE) were used for principal-component analysis (plotPCA) and sample–sample correlation heat-maps (Pearson r, ggplot2 v3.4.4). Differential-expression result tables were exported per comparison, summarised, and filtered lists were subjected to Gene-Ontology over-representation analysis with clusterProfiler (v4.8.3; enrichGO, OrgDb = org.Hs.eg.db v3.17.0, ont = "ALL", padj < 0.05). Transcript-per-million (TPM) values were calculated from raw counts and gene lengths (featureCounts "Length" column) and $\log_2$-transformed for marker-gene heat-maps (pheatmap v1.0.12, viridis palette).

**Deep learning models**

*K-means clustering*

Gene-level TPM values were imported into R (v4.3.2) and filtered to the 10 000 most variable transcripts across all samples (coefficient of variation). Expression values were $\log_2$-transformed and centred before clustering with the base kmeans() function, specifying k = 6, nstart = 25 and the default Euclidean distance. Cluster membership was merged with sample metadata for downstream visualisation of cluster-specific trajectories and Gene-Ontology enrichment.

*Random-forest stage classifier*

For supervised classification, only WT samples were used for training. The TPM matrix was transposed so that rows corresponded to samples and columns to genes; metadata columns (Sample, Genotype, Stage) were removed from the feature set. A factor response vector encoded the four developmental stages (ESC, NE, NSC, NPC). A random-forest model was trained with the randomForest package (v4.7-1.1) using ntree = 500, default mtry, and importance = TRUE. Model performance was inspected via the out-of-bag error rate and a confusion matrix. Variable importance (mean decrease in accuracy) was extracted and plotted for the top 25 genes. The trained classifier was applied to mutant samples (R133C, R168X, R270X) to obtain both hard stage calls and class-probability distributions; results were visualised with ggplot2 (v3.4.4). Mis-classifications were summarized and prediction confidence values (maximum class probability) were compared across genotypes.

*Feed-forward neural-network classifier*

A deep neural network was implemented with the keras R interface (keras v2.13.0; TensorFlow v2.14 backend). WT TPM values were $\log_2(x + 1)$ transformed, gene-wise z-scored (mean subtraction, division by s.d.), and clipped to ±5. The network architecture comprised an input layer matching the number of genes, two hidden dense layers (64 and 32 ReLU units) each followed by 0.30 dropout, and a 4-unit soft-max output layer. Categorical stage labels were one-hot encoded. The model was compiled with categorical cross-entropy loss, the legacy Adam optimizer (learning rate = $5 \times 10^{-4}$), and accuracy as the metric. Training proceeded for up to 200 epochs with batch_size = 4, using a 25 % validation split; early-stopping (patience = 10) and ReduceLROnPlateau (patience = 5) callbacks prevented over-fitting. After convergence, the model weights with the lowest training loss were retained. Mutant samples were pre-processed with the same scaling parameters and predicted to yield both class labels and confidence scores.

**Single-cell RNA Sequencing**

Human ESCs (WT, R133C, R168X and R270X) were harvested at the pluripotent stage, dissociated with Gentle Cell Dissociation (Stem Cell Technologies) and resuspended in ice-cold PBS + 0.04 % BSA. For each genotype, cells from three independent wells were pooled, yielding four fixed suspensions in total. Fixation was performed immediately with the EVERCODE Cell Fixation v2 Kit (Parse Biosciences, #ECF2101) according to the manufacturer's protocol. Cell density and viability were assessed by mixing 10 µL of the suspension with 10 µL 0.4 % Trypan Blue (Thermo Fisher, #T10282) and counting on a Countess 3 Automated Cell Counter; all samples exceeded 90 % viability and were adjusted to $1 \times 10^6$ cells $mL^{-1}$. Two single-cell libraries were constructed from the four fixed samples using the EVERCODE WT Mini v2 Kit (Parse Biosciences, #ECW02115), which employs a split-pool combinatorial indexing workflow comprising four rounds of barcoding followed by cDNA synthesis and amplifications. Final libraries were purified with AMPure XP beads, and double-stranded DNA concentration was measured fluorometrically with a Qubit 4 and the QuantiFluor dsDNA System (Promega, #E2670). Both libraries were sequenced on an Illumina NovaSeq 6000 S4 flow-cell using paired-end 150-bp reads (PE150).

**Single-nucleus RNA sequencing**

Cerebral organoids were harvested on day 90 from three independent cultures per genotype, and all reagents, consumables and centrifuges were pre-cooled to 4 °C. Organoids were transferred to a chilled Dounce containing 700 µL homogenization buffer (NIM1: 250 mM sucrose, 25 mM KCl, 5 mM $MgCl_2$, 10 mM Tris-HCl pH 8, plus 1 mM DTT, 0.40 U $\mu L^{-1}$ RNase-In, 0.20 U $\mu L^{-1}$

Superase-In and 0.1 % Triton X-100) and gently dissociated with 5 loose-pestle and 10 tight-pestle strokes, then adjusted to 1 mL with the same buffer. Lysis efficiency was confirmed by 1:1 Trypan Blue staining, after which lysates were passed through a 40 µm strainer into pre-cooled 15 mL tubes, centrifuged (600 g, 4 min, 4 °C), washed once in PBS containing 0.2 U $\mu L^{-1}$ RNase-In, and re-pelleted under the same conditions. Nuclei were resuspended in 200 µL PBS/RNase-In, re-filtered (40 µm), counted with a Countess 3 Automated Cell Counter (10 µL nuclei + 10 µL 0.4 % Trypan Blue; Thermo Fisher #T10282), and immediately fixed using the Evercode Nuclei Fixation v2 kit (Parse Biosciences, #ECF2103). Eight single-nucleus libraries were then prepared with the EVERCODE™ WT v2 kit (Parse Biosciences, #ECW02135) according to the manufacturer's split-pool protocol. Library yield was quantified with a Qubit fluorometer and QuantiFluor dsDNA system (Promega #E2670), and fragment size distribution verified on an Agilent 2100 Bioanalyzer using the DNA 1000 kit (#5067-1504). Libraries were sequenced on an Illumina NovaSeq 6000 S4 flow-cell using paired-end 150-bp reads (PE150).

**Single-cell and single-nucleus RNA-seq data processing and analysis**

Raw paired-end FASTQ files from hESC single-cell (EVERCODE WT-Mini v2) and 3-month organoid single-nucleus (EVERCODE WT v2) libraries were inspected with FastQC (v0.11.9). Demultiplexing, read trimming, alignment to GRCh38/hg38 and transcript counting were performed with the Parse Biosciences command-line pipeline (commit 2024-02-20); the merge module combined the four sub-libraries per run to produce a gene–cell count matrix (.mtx), feature list and cell-level metadata.

Down-stream analyses were conducted in R (4.3.2) with Seurat (v5.0.1). Matrices were imported with ReadParseBio, and cells with >15,000 detected genes, >100,000 UMIs or >15 % mitochondrial UMIs were removed. Counts were log-normalised (NormalizeData, scale.factor = 10,000), 2,000 variable features were selected (FindVariableFeatures, method = "vst"), and expression values were centered and scaled (ScaleData). Dimensionality reduction used PCA (RunPCA); the first 30 principal components (PCs) were retained based on the elbow plot. A shared-nearest-neighbour graph was constructed (FindNeighbors, dims = 1:30) and Louvain clusters identified at resolution = 0.30 (FindClusters). Clusters were reordered by hierarchical tree building (BuildClusterTree) and visualised with UMAP (RunUMAP, dims = 1:30). Cluster markers were identified with FindAllMarkers (min.pct = 0.25, logfc.threshold = 0.25) and visualised by violin, dot, and feature plots.

**Quantification and statistical analysis**

All statistical procedures, n values, definitions of center, dispersion metrics, and exact *P* or adjusted *P* values are reported in the corresponding figure legends, main text, or Method subsections. A consolidated summary of analytical strategies is provided below.

| Experiment | Software | Statistical test or model | Definition of *n* | Measures of center ± dispersion | Multiple-test correction | Location |
|---|---|---|---|---|---|---|

| | | | | | | |
|---|---|---|---|---|---|---|
| **RT-qPCR (fig. 3D and 4B)** | Prism 10 (GraphPad) | One-way ANOVA with Dunnett post-hoc (mutant vs WT) | Independent iPSC or hESC cultures | Mean ± s.e.m. | — | Fig. 3D and 4B legend |
| **Cell-growth (fig. 3H)** | Incucyte S3 software + Prism 10 | Two-way repeated-measures ANOVA | Colonies tracked over time (9 fields × 3 wells × 3 experiments per genotype) | Mean ± s.e.m. | — | Fig. 3H legend |
| **Cell-cycle phase counts (fig. 3G)** | Seurat 5.0.1 | Pearson $\chi^2$ goodness-of-fit vs WT | Single cells passing QC (exact cell counts in legend) | Proportion (% of cells) | — | Fig. 3F–G legend |
| **Bulk rna-seq DE (figs. 1, 4)** | DESeq2 1.40.2 | Wald test (mutant vs WT) | Independent differentiations (3 per genotype & stage) | $\log_2$ fold-change | Benjamini–Hochberg padj < 0.01 & | $\log_2$FC |
| **GO enrichment (fig. 1C, 2C)** | clusterProfiler 4.8.3 | Hypergeometric test | Gene universe = all expressed genes | GeneRatio, $-\log_{10}$ padj | Benjamini–Hochberg padj < 0.05 | Methods → "Bulk RNA-seq analysis" |
| **Machine learning (fig. 2)** | R 4.3.2: stats::kmeans, randomForest 4.7-1.1, keras 2.13/TensorFlow 2.14 | — | See below | OOB error, prediction probability | 5-fold CV for RF; 25 % validation split for NN | Fig. 2 legends |

*Randomization, blinding, and sample-size considerations.*

hESC and iPSC lines were plated and differentiated in parallel under identical conditions; wells were assigned randomly to imaging positions in Incucyte assays, and image acquisition/analysis were automated. Organoid batches were generated from separate vials per genotype to preserve biological replication. Investigators were not blinded to genotype during cell culture but downstream bioinformatic pipelines were scripted and executed without manual intervention. For exploratory *in vitro* systems we did not perform a priori power calculations; instead, sample sizes (three independent differentiations per condition, ≥2 900 single cells per scRNA-seq sample, three organoids per genotype for snRNA-seq) reflect field standards and our prior experience achieving reproducible effect sizes.

*Inclusion and exclusion criteria.*

For RT-qPCR and growth assays all biological replicates were included. sc/snRNA-seq data were filtered to retain nuclei with <15 % mitochondrial RNA and 500 ≤ nFeature_RNA ≤ 15 000; doublets detected by index hopping were excluded. Bulk RNA-seq genes with ≤1 read across all samples were removed prior to DESeq2.

*Definition of significance.*

Unless otherwise specified, $P < 0.05$ (after correction where applicable) was considered significant. Exact $P$, padj, or FDR values are provided in figure panels or legends.
